# Fully Implanted Prostheses for Musculoskeletal Limb Reconstruction after Amputation: An *In Vivo* Feasibility Study

**DOI:** 10.1101/2020.07.02.184994

**Authors:** Patrick T. Hall, Samantha Z. Bratcher, Caleb Stubbs, Rebecca E. Rifkin, Remi M. Grzeskowiak, Bryce J. Burton, Cheryl B. Greenacre, Stacy M. Stephenson, David E. Anderson, Dustin L. Crouch

**Affiliations:** University of Tennessee, Knoxville – Knoxville, TN – Department of Mechanical, Aerospace, and Biomedical Engineering, 1512 Middle Dr, Knoxville, TN 37966; University of Tennessee College of Veterinary Medicine – Knoxville, TN – Department of Large Animal Clinical Sciences; University of Tennessee College of Veterinary Medicine – Knoxville, TN – Office of Laboratory Animal Care; University of Tennessee College of Veterinary Medicine – Knoxville, TN – Department of Small Animal Clinical Sciences; University of Tennessee Medical Center – Knoxville, TN

**Keywords:** Orthopedic, Vascularization, Subdermal, Endoprosthesis, Osseointegration, Animal Model

## Abstract

Previous prostheses for replacing a missing limb following amputation must be worn externally on the body. This limits the extent to which prostheses could physically interface with biological tissues, such as muscles, to enhance functional recovery. The objectives of our study were to (1) test the feasibility of implanting a limb prosthesis, or endoprosthesis, entirely within living skin at the distal end of a residual limb, and (2) identify effective surgical and post-surgical care approaches for implanting endoprostheses in a rabbit model of hindlimb amputation. We iteratively designed, fabricated, and implanted unjointed endoprosthesis prototypes in six New Zealand White rabbits following amputation. In the first three rabbits, the skin failed to heal due to dehiscence along the sutured incision. The skin of the final three subsequent rabbits successfully healed over the endoprotheses. Factors that contributed to successful outcomes included modifying the surgical incision to preserve vasculature; increasing the radii size on the endoprostheses to reduce skin stress; collecting radiographs pre-surgery to match the bone pin size to the medullary canal size; and ensuring post-operative bandage integrity. These results will support future work to test jointed endoprostheses that can be attached to muscles.

## Introduction

A major goal of limb prostheses is to return intuitive control and realistic sensation of movements associated with the lost limb following amputation. Such sensorimotor function is especially important for upper^15, 24, 30^ limb amputees since it enables the closed-loop motor control needed to perform manual, dexterous tasks. This goal has hitherto been pursued using external (i.e. worn outside of the body) prostheses. Most external prostheses incorporate electromechanical hardware (e.g. motors, microprocessors, sensors, electrodes)^17, 23^ and algorithms that decode users’ movement intent from electromyograms^5, 14^ and stimulate nerves to provide sensory feedback. Electromechanical systems do not yet accurately replicate sensorimotor physiology and, thus, likely introduce errors in decoding intended movement patterns, generating prosthesis movements, and delivering proper sensory feedback that directly correlates with the movements. The sensorimotor limitations of external myoelectric prostheses are a major reason why many are eventually abandoned^2^.

One potential solution to overcome sensorimotor limitations of external prostheses is to physically attach prostheses to muscles. This is because muscles contribute to both movement generation and sensation, via mechanoreceptors, in the biological limb. There is strong evidence that muscles in the residual limb retain their sensorimotor functions after amputation^4, 5, 11, 28^. Since all current prostheses are worn externally, the previous attachment approach required transferring muscle forces through skin using a procedure called cineplasty^3, 30^. Reports note anecdotally that cineplasty enabled exquisite control and sensation of prosthesis movements^8, 30^. However, cineplasty has several limitations in function, comfort, and appearance that are directly related to the need to transfer muscle forces through skin. Given these limitations, cineplasty has not been widely adopted.

Our proposed novel approach to better facilitate physical muscle-prosthesis attachment is to implant prostheses completely within skin. With an implanted prosthesis, or *endo*prosthesis, the residual muscles could be attached in a more cosmetic and anatomically realistic way that would overcome the limitations of cineplasty. Like other common orthopedic implants such as joint replacements^10, 13^, endoprostheses would replace part of the musculoskeletal structure of the missing limb. The proposed endoprosthesis differs from previous percutaneous osseointegrated prostheses, which protrude through skin to provide an anchor point for externally worn limb prostheses.

The objectives of our proof-of-concept study were to (1) test the feasibility of implanting a limb prosthesis, or endoprosthesis, entirely within living skin at the distal end of a residual limb, and (2) identify effective surgical and post-surgical care approaches for implanting endoprosthesis prototypes in a rabbit model of hindlimb amputation. We iteratively implanted an unjointed endoprosthesis prototype *in vivo* in six New Zealand White rabbits with transtibial (i.e. below-knee) amputation. We refined the surgery, post-operative care, and endoprosthesis design between surgery iterations. Our criteria for a successful proof of concept was that at least two rabbits with the endoprosthesis could recover enough post-surgery to forego bandages and medications.

## Materials and Methods

### Summary

This study was approved by the University of Tennessee, Knoxville Institutional Animal Care and Use Committee. We used six male New Zealand White rabbits (R1-R6; average pre-surgery weight = 2.92 kg, average age = 18 weeks), since the rabbit is a common orthopedic model and large enough for testing physical endoprosthesis prototypes compared to other smaller mammals. We used an iterative study design; in the same survival surgery and in one rabbit at a time (from R1 to R6), we amputated the hind limb below the knee and implanted the stem endoprosthesis. We monitored the rabbits during post-surgical recovery, qualitatively noting any adverse events that negatively affected the surgical outcome. We reviewed the observations as a team and adapted our surgery or device as necessary to address the problems in the subsequent iterations. Rabbits were euthanized either at a humane endpoint or after post-surgical recovery (at around 60 days post-surgery), whichever was sooner. The rabbits were housed with at least one companion rabbit in adjacent cages. Enrichment and positive human interaction were given to the rabbits daily.

### Stem Endoprosthesis

The stem endoprosthesis was comprised of three parts: a metal segment, a modified intramedullary bone pin, and an over-molded silicone sleeve (Fig. 1). The metal segment was designed in Solidworks (Dassault Systemes, France) and 3D-printed in 316 stainless steel. We drilled and tapped a hole on the flat proximal surface of the stem with a reverse thread for later integration with the bone pin. Four of the stems (R3-R6) had sites for muscle attachment to stabilize the stem in the bone and mimic future muscle attachment to a muscle-driven endoprosthesis; the muscle attachment sites were added at will and not in response to an adverse surgical outcome. Commercially available intramedullary bone pins (IMEX Veterinary, Inc.), which we used to anchor the stems to the tibia bone, were cut to a shorter length and tapped with additional threads to screw into the metal segment. A range of pin sizes were available (Table 1). The metal segment was over-molded with an approximately 2-mm-thick coating of biocompatible silicone of hardness 40 shore A (BIO LSR M140, Elkem Silicones) using a custom, 3D-printed 316 stainless steel mold. Once the three parts were assembled, the stems were sterilized using ethylene oxide gas.

**Table 1.** Available intramedullary bone pin sizes.

| Imex Bone Pin Diameter | Outer Thread Diameter |
| --- | --- |
| 1.6 mm | 1.9 mm |
| 2.0 mm | 2.3 mm |
| 2.0 mm | 2.5 mm |
| 2.4 mm | 3.2 mm |
| 2.8 mm | 3.5 mm |

**Figure 1.**
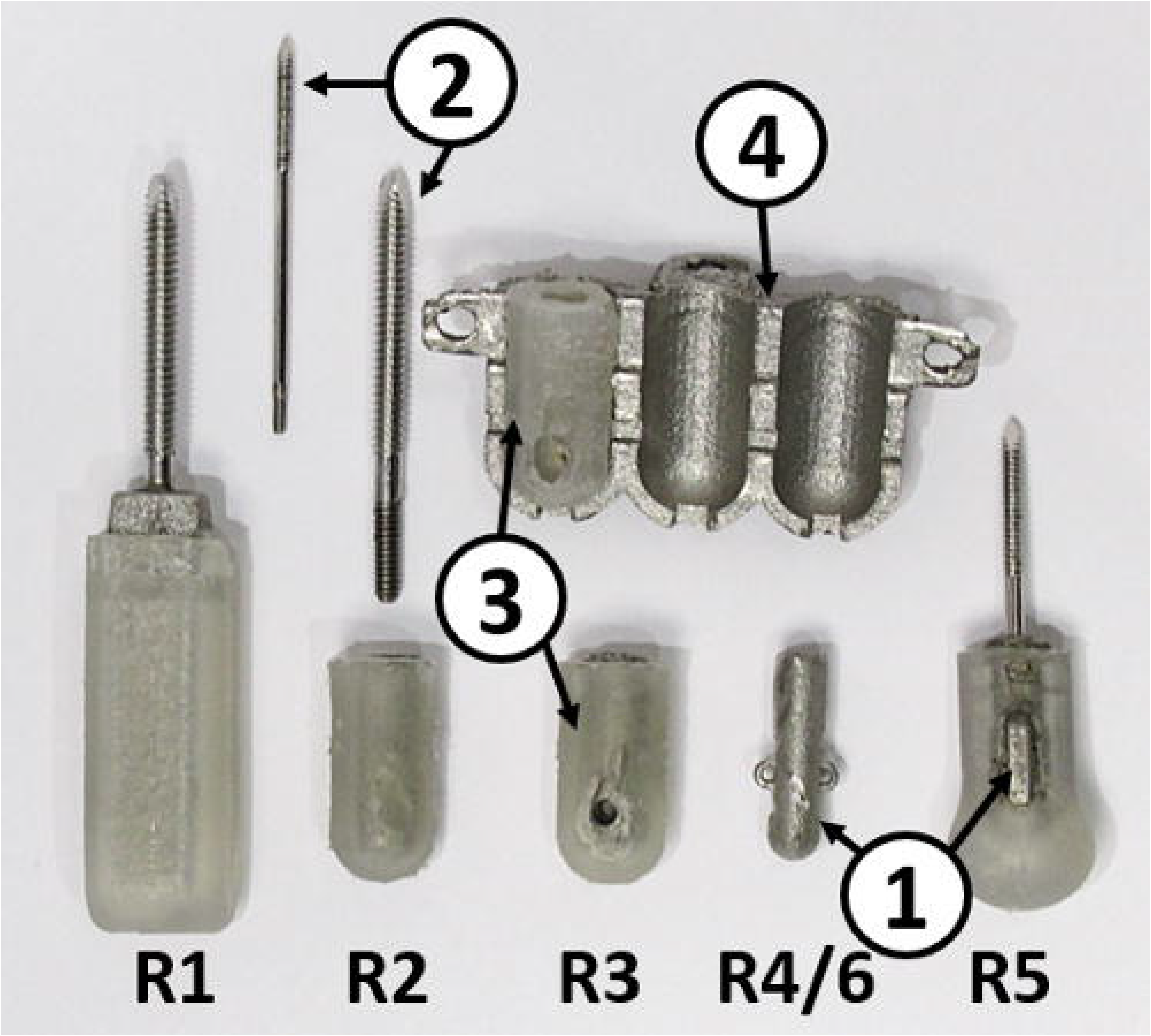
Stem endoprostheses used during surgery. Corresponding animal number noted below each device. (1) 3-D printed 316L stainless-steel stem. (2) modified threaded intramedullary bone pin. (3) biocompatible, over-molded silicone sleeve. (4) 3-D printed stainless-steel mold used to over-mold silicone onto stem.

### Pre-Surgery Bone Geometry Estimation

We amputated the hindlimb approximately 4 cm from the distal end of the tibia. This allowed us to screw the bone pin into the approximate mid-diaphyseal area of the tibia where the medullary canal is narrowest. We approximated the canal diameter from either micro-computed tomography images from another rabbit of similar size (R1) or pre-surgery radiography images of the operated rabbits (R2-R6).

### Surgical Technique

The general surgical procedure for each rabbit was as follows, although we modified some steps in each surgery iteration (see Results) based on the surgical outcomes of preceding iterations. The rabbits were given a pre-emptive analgesic, either buprenorphine (0.03-0.05 mg/kg) or hydromorphone (0.2 mg/kg), and induced into general anesthesia with either a solution of ketamine (30-40 mg/kg) and xylazine (3-7 mg/kg) or midazolam (0.75-1 mg/kg). Anesthesia was maintained with 3-5% isoflurane gas. We removed the hair from the operated limb with electric clippers and depilatory cream (Nair Hair Remover Cream, Church & Dwight Co., Ewing Township, NJ). We positioned the rabbit in right lateral recumbency with the left leg suspended, and aseptically prepared the limb with chlorohexidine, betadine, and 70% isopropyl alcohol. The surgeons (Anderson, Rifkin, and Grzeskowiak) made an incision on the hindlimb preserving enough skin and soft tissue to create a skin flap to cover the implanted stem. The surgeons then used an osteotomy saw blade to cut the bone in the distal tibial diaphysis. The bone pin of the stem was then screwed into the exposed tibial medullary canal. For the stems for R3-R6, which had sites for muscle attachments, the surgeons used a 5-0 synthetic absorbable monofilament suture (PDS) to anchor the gastrocnemius tendon and the tibialis cranialis insertion tendon to the stem with a locking loop. The skin flap created during the incision was wrapped across the distal end of the stem and sutured closed in a continuous subcuticular pattern using a 4-0 synthetic absorbable monofilament suture. In R2-R6, the closure was reinforced with a 2-0 synthetic absorbable monofilament applied in an interrupted cruciate pattern. Liquid topical tissue adhesive (3M Vetbond Tissue Adhesive) was placed over the external surface of the incision line.

### Post-Surgical Care

Silver sulfadiazine topical cream was applied over the incision to prevent infection, but immediately post-operatively and at every bandage change. The limb was bandaged using, from inner to outer layers, non-adherent dressing, undercast padding, elastic bandage wrap, and elastic tape (ELASTIKON, Johnson & Johnson) to protect the incision site. We changed the bandage at least once every three days and monitored skin integrity and incision healing. We administered an analgesic (buprenorphine 0.03 mg/kg) subcutaneously every 6 hours for at least 72 hours post-surgery, as well as antibiotics every 12 hours for at least 7 days (enrofloxacin 5 mg/kg diluted) and an anti-inflammatory drug every 24 hours for at least 7 days (meloxicam; 0.6 mg/kg) subcutaneously for at least 7 days post-surgery.

### Imaging

We acquired radiographs of the operated limb postoperatively and approximately every two weeks post-surgery to monitor the bone pin alignment, stem position, and bone.

### Weight Bearing Analysis

After the skin healed and fully enclosed the endoprosthesis in R4 and R6, we removed the bandage. To determine the extent to which the rabbit was willing to apply pressure to the endoprosthesis limb, we conducted a testing session to measure ground-limb pressures. The rabbit was guided onto a pressure mat (Tekscan Very HR Walkway 4) and we measured the pressure of both the operated and intact contralateral limbs simultaneously over an 8-second period while the rabbit was standing still. We compared, between limbs, the magnitude of both vertical pressure and total vertical force applied by each limb.

## Results

### Bone-Device Interface

When inserting the bone pin in the tibia of R1, the dorsal tibial cortex fractured, requiring a more proximal amputation to remove the fractured bone. Since the amputation was performed more proximally than intended, the threads of the bone pin were aligned with the wider area of the tibia medullary canal, creating a loose fit between the bone and bone pin (Fig. 2). Therefore, bone cement (polymethyl methacrylate) was placed into the medullary canal to secure the bone pin. The tibia fractured in R1 because the bone pin diameter, estimated from micro-computed tomography images from another rabbit of comparable size, was too large. We also had only one bone pin size available for R1, which prevented the surgeons from switching pins during surgery in case of a size mismatch. Therefore, for the next five surgeries (R2-R6), the tibial medullary canal measurements were determined from *in vivo* radiographs of each rabbit taken 1-2 weeks prior to surgery. We selected the diameter of the bone pin to be the same as or slightly less than the medullary canal diameter so that the pin would fit snugly in the canal without fracturing the tibia. Additionally, three different bone pin size options were available if determined to be needed intraoperatively: the pin with a thread diameter nearest the smallest medullary canal diameter measured from radiographs, the next larger pin, and the next smaller pin. Table 1 outlines the available pin sizes from which we selected. Using this approach, for R2-R6, we experienced no complications with the bone pin fit and achieved a snug fit without need for bone cement. The endoprosthesis or bone pin did not appear to move based on the post-surgical radiographs (Fig. 2). Post-surgical radiographs indicated proliferative bone ongrowth over the cranial aspect of the device as early as 12 days after surgery in all rabbits (Fig. 2). However, this bone growth was visibly smaller in R4-R6, which had the muscle attachments.

**Figure 2.**
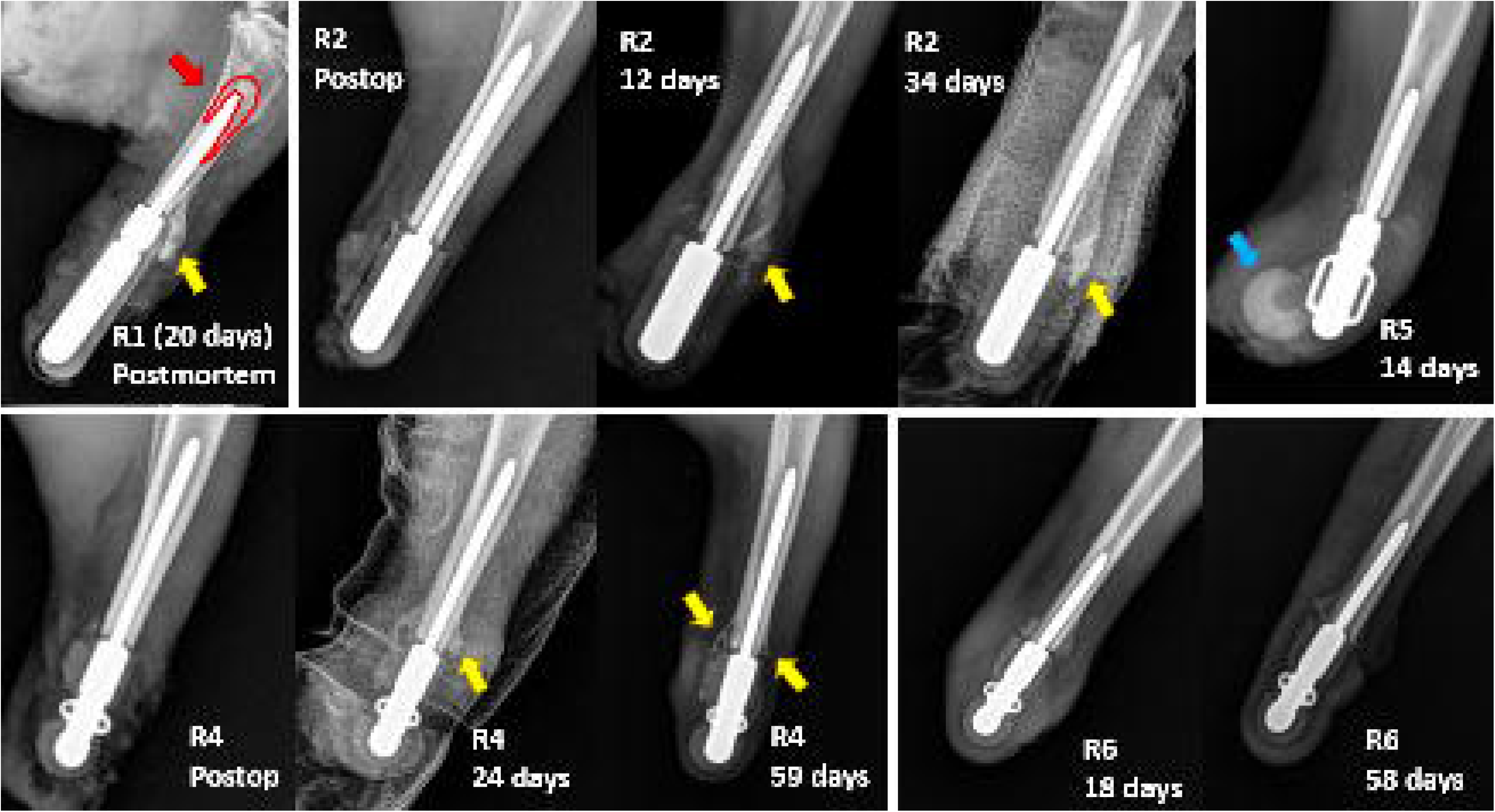
Radiographs of stems in situ. Top Left: R1 postmortem 3-weeks post-surgery showing bone cement (red outline and arrow) and proliferative bone ongrowth indicated by yellow arrow. Top Middle: Progression of R2 over 5 weeks showing development of bone ongrowth (yellow arrow). Bottom Left: Progression of stem in R4 showing development of bone ongrowth (yellow arrow). Top Right: X-ray of R5 two weeks post-surgery showing where the silicone slipped off the end of

### Skin Closure

For R1, we used a cranial circumferential incision to create a caudal skin flap and cranial suture line. This preserved the calcaneal fat pad to absorb the pressure applied to the skin when the rabbit bore weight on the residual limb. At 3 days post-surgery, the skin surrounding part of the implant was dark, indicative of ischemia (Fig. 3). A portion of the incision line dehisced by 10 days. At 3 weeks, the skin at the distal end of the residual limb broke off at the edge of the ischemic area, exposing the stem and requiring euthanasia of the rabbit. The skin complications in R1 were potentially due to the stem geometry and incision technique. The stem for R1 was relatively long and had corners with relatively small radii, which could have created areas of high mechanical stress in the skin. In subsequent surgeries, we made the stem shorter and increased the radius of the distal end of the stem, Additionally, for R2 and R3, we reversed the incision to create a caudal skin closure to prevent weight bearing and loading on the incision line.

**Figure 3.**
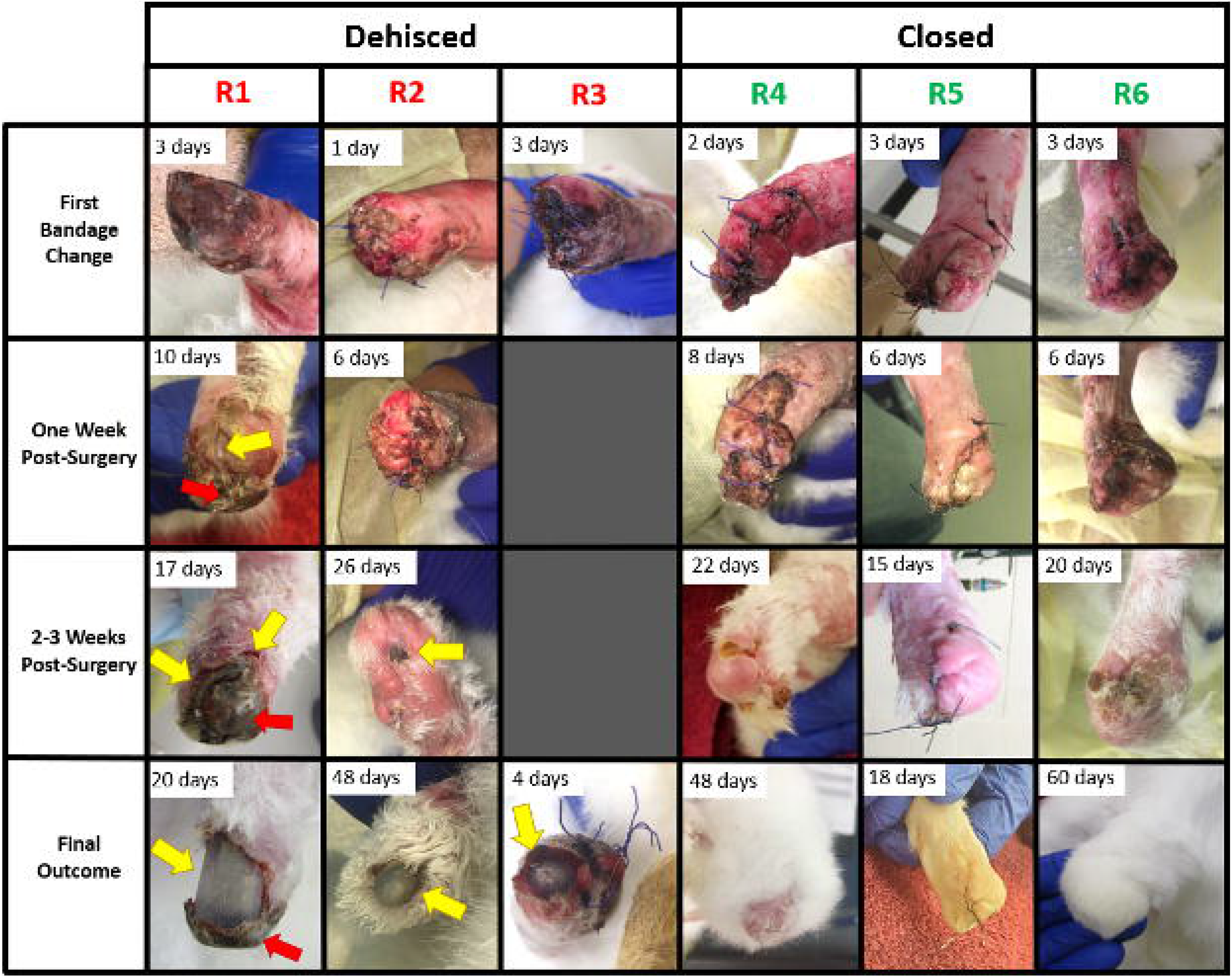
A timeline displaying how the skin of the operated limb for each rabbit changed over time. Pictures were taken at every bandage change, and progress noted at various timepoints. The number of days after surgery the picture was taken is in the top left corner of each cell. Yellow arrows indicate dehiscence in the skin. Red arrows indicate ischemic skin locations. The animal numbers indicate the order of surgeries (i.e. R1 was performed first, then R2, etc.).

R2 showed no sign of ischemia (Fig. 3). However, at 3 weeks post-surgery, the skin had failed to completely heal and a small dehiscence (~2 mm diameter) had formed along the incision line.

The dehiscence enlarged by about 7 weeks, exposing the stem and requiring euthanasia of the rabbit. At 4 days post-surgery, R3 had removed the bandage and appeared to have chewed on the sutures and stem, causing the incision line to dehisce. Attempts to close the incision failed, so the rabbit was euthanized.

Although we had only observed ischemia in R1, we were concerned that the circumferential incision traditionally used during amputation could disrupt the microvasculature arising from the femoral artery that runs caudally along the hindlimb^9^. Therefore, starting with R4, we altered the surgical approach to a cranial linear incision to preserve the vasculature (Fig. 4). This technique is similar to one used in human knee arthroplasty^25^. By as late as 22 days post-surgery in R4-R6, the skin was healthy and the incision line healed (Fig. 3). After determining successful healing of the residual limb, we permanently removed the bandage. In R4 there were some areas of skin over the distal end of the stem that appeared bruised, presumably from impacts against the ground or cage bottom, but the skin remained closed over the stem. Pressure analysis of R4 and R6 showed that the residual limb experiences pressures of up to 0.92 kg/cm^2^, which was 3.28 times that of the maximum intact contralateral limb under general weight bearing (Table 2). On average, the residual limb experienced about 2.6 times greater pressure than the intact contralateral limb over an 8 second timeframe. This is due to the smaller surface contact area in the residual limb. Despite high pressure, the rabbits were willing to place the equivalent of about 25% body weight on the endoprosthesis.

**Table 2.**
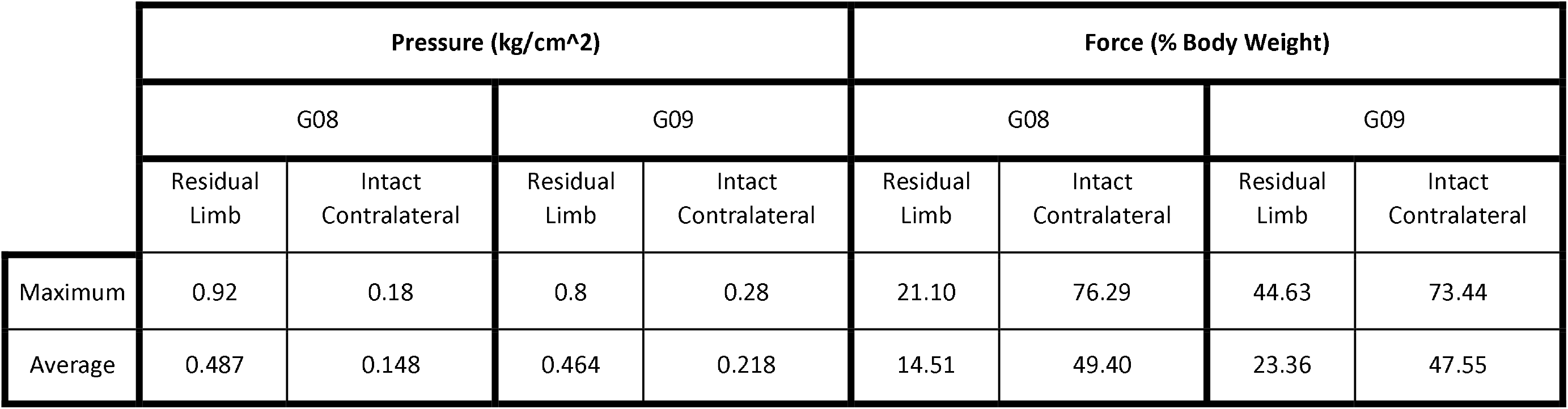
Pressure analysis of rabbits when bearing weight on all four legs while standing.

|  | Pressure (kg/cm<sup>2</sup>) |  |  |  | Force (% Body Weight) |  |  |  |
| --- | --- | --- | --- | --- | --- | --- | --- | --- |
|  | G08 |  | G09 |  | G08 |  | G09 |  |
|  | Residual Limb | Intact Contralateral | Residual Limb | Intact Contralateral | Residual Limb | Intact Contralateral | Residual Limb | Intact Contralateral |
| Maximum | 0.92 | 0.18 | 0.8 | 0.28 | 21.10 | 76.29 | 44.63 | 73.44 |
| Average | 0.487 | 0.148 | 0.464 | 0.218 | 14.51 | 49.40 | 23.36 | 47.55 |

**Figure 4.**
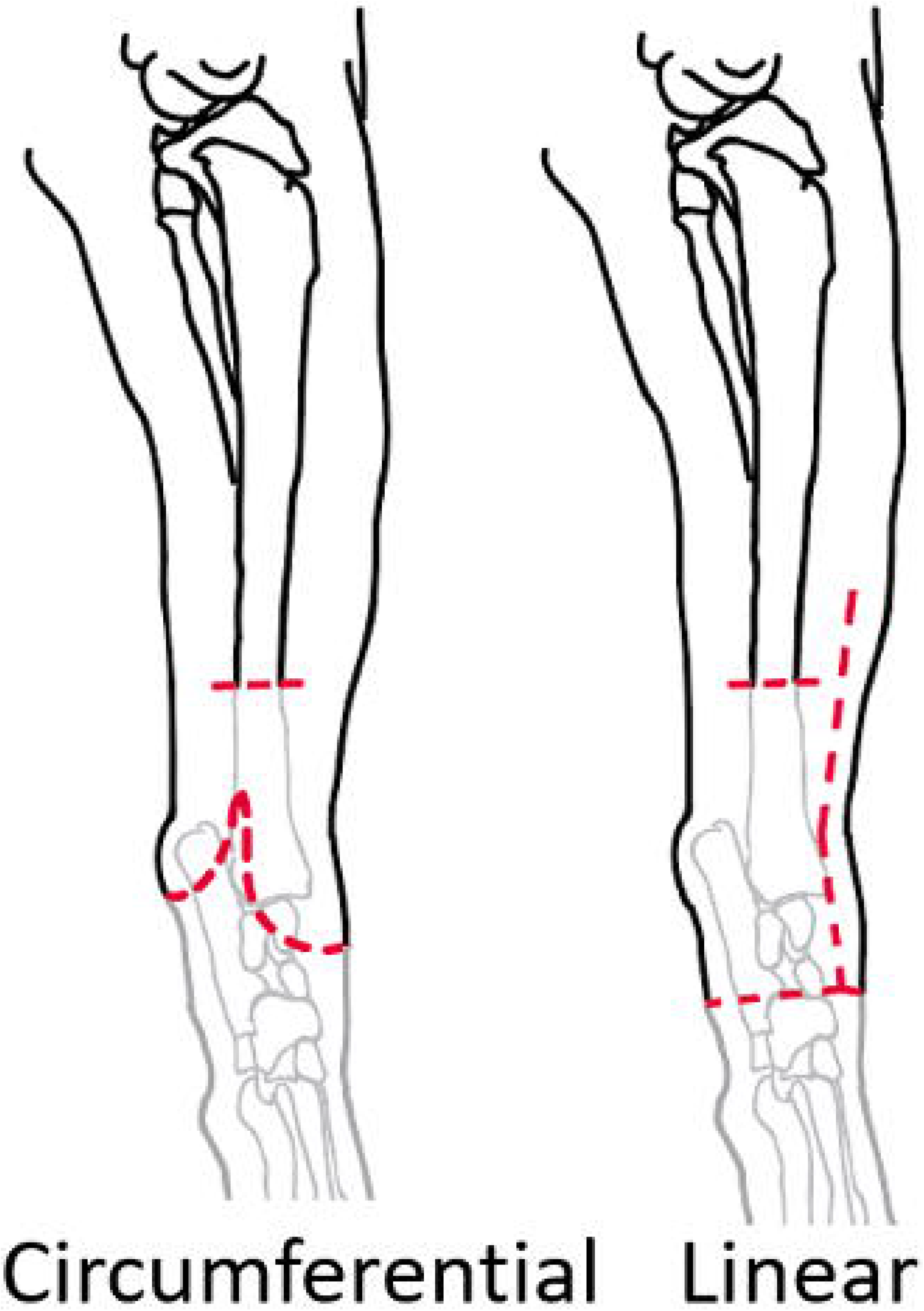
Representation of the change in the incision approach that occurred between R3 and R4. We switched to a linear incision to preserve the vasculature of the skin flap provided through the femoral artery. The tibia was amputated at approximately the same location for both incision approaches.

### Bandaging and Recovery

Rabbits R1-R3 were able to remove the bandages on their own, potentially exposing the operated limb to trauma and infection. The bandage removal may have contributed to dehiscence in R1 and R2. There were clear indications that R3 chewed on the stem, suggesting that the rabbit may have also caused the dehiscence after removing the bandage. We took several steps to try to prevent the rabbits from removing the bandages. We monitored the bandages more closely, at least twice per day, for two weeks post-surgery. If we noticed that the bandage was damaged or beginning to slip off, we applied additional elastic tape to reinforce the bandage. For R4-R6, we preemptively applied an extra bandage layer, which drastically reduced the number of times that the rabbits were able to remove the bandage.

### Device Design

As noted above, we modified the implant design after R1 by making the endoprosthesis shorter (20mm vs 40mm) and with larger radii to improve the chance of successful skin closure and reduce stress concentrations on the skin. To further decrease pressure on the skin in R5, we increased the radius of the silicone sleeve tip from 4.5mm to 10mm by increasing the thickness of the silicone cover without changing the design of the metal segment. We performed the same surgical technique as R4 to implant the stem in R5. However, radiographs of R5 at 14 days post-surgery revealed that the larger silicone sleeve had slipped off the tip of the metal segment (Fig. 2). We were able to realign the silicone temporarily, but it would not stay in place. Therefore, given the potential high risk of skin trauma against the exposed metal segment, we euthanized R5 once the bandaged skin incision had healed and fully closed (18 days post-surgery).

## Discussion

In this first-of-its-kind study, we demonstrated convincingly in an *in vivo* model that it is feasible to fully enclose an endoprosthesis in living skin at the distal end of a residual limb. That all three final surgeries resulted in a successful shows the repeatability of our approach. Closing living skin over an endoprosthesis is a major challenge. This is partly because the skin at the distal end of the rabbit hindlimb is very thin, making it difficult to achieve suitable apposition of the wound edges along the suture line to promote wound healing. Additionally the suture line closing the wound lies directly over the synthetic endoprosthesis, which could interfere with wound healing^27^. As with all wounds, the suture itself can damage tissue and, if tied too tight, crush cells or occlude blood flow to the wound^22^. Despite these challenges, our approach was sufficient to achieve wound closure by about 3 weeks post-surgery.

Preserving the blood supply to the skin is critical for wound healing and for maintaining long-term skin health. Along the length of an intact limb, branches from deep, central vessels extend superficially into the skin’s vascular network. In rabbits, the terminal arterial supply is extremely fragile and susceptible to disruption. Additionally, the skin covering the endoprosthesis cannot receive blood flow from such deep branches and, thus, must rely only its own vascular network. The central vessels in the rabbit hindlimb, the femoral and saphenous arteries, run along the caudal aspect of the hindlimb. During the first three surgeries, we used a circumferential incision with proximal reflection because it is the more traditional approach for limb amputation and closure of the skin around the residual limb. However, the circumferential incision, especially in R1, may have severed branches derived from the femoral or saphenous arteries and decreased the blood flow to the incision site. Thus, for rabbits R4-R6, we changed to a linear incision, an approach used in other orthopedic procedures^25^. Though the wound closed in all three rabbits after changing the incision, additional studies would be needed to confirm whether the change in surgical approach was the reason for improved outcomes.

Endoprostheses may place relatively high stress on the suture line and overlying skin since, unlike traditional orthopedic implants, they directly contact the skin. It has been shown in both pig^16^ and rabbit^29^ models that some mechanical loading along the skin surface and directly across a wound encourages healing. However, the skin over an endoprosthesis would also experience pressure on the suture line due to ground contact forces, which could damage the suture line and cause dehiscence. We took two steps to reduce the risk of skin trauma with the endoprosthesis. First, we applied the padded bandage over the limb for about 3 weeks after surgery, which reduce the pressure applied to the suture line as it healed. Second, we covered the endoprostheses with a compliant material (silicone), which would reduce skin pressure when external loads are applied. Additional studies are needed to determine the resilience of the skin covering the endoprosthesis to trauma under typical and severe biomechanical loading conditions.

Silicone, the material selected as the compliant coating for our endoprosthesis prototypes, has been used in some orthopedic devices to, for example, prevent soft tissue adhesions that would interfere with the implant’s function^21^. Silicone is also common in other subdermal implants used in cosmetic surgery^7^. A potential drawback of coating the metal segment in silicone is that it could increase the risk of infection. Advanced multi-step sterilization procedures or incorporation of anti-bacterial coatings may be needed to prevent infection with long-term use of silicone-coated devices. Alternatively, biomaterials could be used to provide mechanical compliance and other benefits. For example, integration of a collagen layer over a tracheal prostheses improved the rate of epithelization over the prosthesis^26^. Biomaterial coatings over an endoprosthesis could promote wound healing and make the skin over the endoprosthesis more resilient to trauma.

Like many other orthopedic implants, such as joint replacements, our endoprosthesis was osseointegrated in the medullary canal of a long bone. Osseointegration is also becoming more widely used to attach external limb prostheses to a residual limb^12, 19^ since it overcomes problems of external prosthesis sockets such as skin breakdown and discomfort. In our proof-of-concept study, for convenience, we attached the endoprostheses to bone using off-the-shelf stainless-steel bone pins with machined threads. This osseointegration approach is not as sophisticated and effective as the clinical and state-of-the-art devices that incorporate, for example, porous metal frameworks^20^ and osseoinductive coatings^1^. Such features encourage bone cells to adhere to or grow within the device so that it is more structurally stable relative to the bone^6, 18^. Of course, in future studies, our endoprosthesis could leverage state-of-the-art design features to achieve more effective long-term osseointegration.

We are unaware of previous reports that show similar bone ongrowth over the exterior of an osseointegrated implant as we observed in the post-surgery radiographs (Fig. 2). The bone ongrowth is likely a biological response by the bone to achieve better mechanical fixation and may have been further stimulated by external loading from ground-limb contact. Since the bone growth occurs cranially and caudally, it could potentially interfere with muscles attached to the endoprosthesis by protruding into the muscle’s path. However, when we attached muscles to the endoprosthesis in R3-R6, we qualitatively observed less proliferative bone ongrowth, supporting the notion that the muscle attachment improved mechanical stability and fixation of the endoprosthesis in the bone. We plan to conduct histology to better understand the structure of the bone ongrowth tissue. Quantifying the extent of bone ongrowth and evaluating its effect on the performance of muscle-driven endoprostheses will also be part of our future research.

The unjointed endoprosthesis presented in this study is a first step toward our larger goal to develop jointed endoprostheses that are physically attached to muscles. Muscle-driven endoprostheses could potentially restore realistic movement control and proprioception in people with amputation far beyond the level enabled by current external prostheses that remain physically detached from muscles. Additional benefits of endoprostheses include a more natural cosmetic appearance; fewer skin problems associated with external prostheses; lower prosthesis weight compared to electromechanical prostheses that require motors and batteries; and more convenience without the need to recharge or replace batteries. The potential benefits of endoprostheses would radically improve function and quality of life of people with limb amputation and other major musculoskeletal defects.

Our study had several limitations. First, we used the native skin of a healthy rabbit to enclose the endoprosthesis. When implanting an endoprosthesis in a person who has already undergone amputation, enough native skin to cover the implant would not be present. Thus, future research should investigate approaches, such as tissue expansion or skin grafting, to increase the amount of skin available to cover an endoprosthesis. Second, we had a small sample size (n=6), which was sufficient for our proof-of-concept study, though larger sample sizes will be needed to evaluate endoprostheses while accounting for inter-specimen variation. Third, the duration of study was relatively short. The rabbits with successful outcomes were kept alive for up to 60 days, which was enough to achieve our study objective but too short to determine long-term effects on tissues. Fourth, the endoprosthesis did not incorporate many state-of-the-art features for osseointegration, as described above, or materials that are common in orthopedic implants. For example, we used 316L stainless steel rather than titanium, which is traditionally used in orthopedic implants, because of our ability to 3-D print stainless steel in-house to quickly and affordably iterate on our endoprosthesis design. However, state-of-the-art materials and features could easily be incorporated, which we expect would only enhance the biocompatibility and performance of endoprostheses.

In conclusion, we showed that it is feasible to fully enclose an endoprosthesis in living skin at the distal end of a residual limb. We identified several critical factors that need to be considered when implanting an end-of-limb endoprosthesis, such as preserving vascularization, protecting the limb, and limiting the pressure applied to the skin. Our next steps include (1) comprehensively evaluating the interfacing bone and skin tissues using imaging and histology, and (2) testing progressively larger implants, with the ultimate goal of implanting a jointed, muscle-driven endoprosthesis prototype to replace the foot and ankle of the rabbit hindlimb. Our repeatable success in implanting unjointed endoprostheses is a promising early step toward realizing our revolutionary and transformational muscle-driven endoprosthesis concept.

## Acknowledgements

The authors thank Elizabeth Croy for her assistance with surgeries and radiography; Dr. Lori Cole and Chris Carter for veterinary care provided for the rabbits in this study; the Office of Laboratory Animal Care and Animal Housing Facility staffs at the University of Tennessee, Knoxville for animal care assistance; Dr. William Hamel for use of the stainless-steel 3-D printer; Dr. Brett Compton for providing silicone preparation equipment; and Danny Graham for his assistance in machining parts for the endoprostheses. Research reported in this publication was supported by (1) the Eunice Kennedy Shiver National Institute of Child Health & Human Development of the National Institutes of Health under Award Number K12HD073945, (2) a seed grant from the University of Tennessee Office of Research and Engagement, and (3) the University of Tennessee Department of Mechanical, Aerospace and Biomedical Engineering start-up funds.

